# Comparative essentialome analysis of six *Pectobacteriaceae* strains using the TNSEEK pipeline identifies conserved and strain-specific fitness determinants

**DOI:** 10.64898/2025.12.09.693261

**Authors:** Julie Baltenneck, Loïc Couderc, Jacques Pédron, Guillemette Marot, Areski Flissi, Hélène Touzet, Erwan Gueguen, Marie-Anne Barny, Guy Condemine

**Author notes:** Co first authors listed in alphabetical order (Julie Baltenneck, Loïc Couderc, Jacques Pédron). Co last authors listed on the basis of seniority (Erwan Gueguen, Marie-Anne Barny, Guy Condemine).

## Abstract

Transposon sequencing (Tn-seq) is a powerful technique for defining the essential genes required for bacterial survival. However, gene essentiality can vary significantly across taxonomic levels, and comparing large Tn-seq datasets from multiple strains presents considerable analytical challenges. To address this, we developed TNSEEK, a fully automated bioinformatics pipeline for the systematic and comparative analysis of transposon sequencing experiments. We applied TNSEEK to analyze six Soft Rot *Pectobacteriaceae* (SRP) strains, encompassing species from the *Dickeya* and *Pectobacterium* genera, grown in a rich medium. This approach identified a core essentialome of 225 genes, primarily involved in fundamental cellular maintenance, conserved across all six strains, a set comparable in size to that of the broader *Enterobacteriaceae* family. Only a few genus-specific essential genes were found highlighting interesting distinct metabolic capabilities between *Dickeya* and *Pectobacterium* genera. In striking contrast, we discovered a large variable essentialome comprising 181 strain-specific genes, many of which of unknown function. A portion of these strain-specific essential genes are components of defense systems and prophage genomic regions. The unexpected essentiality of these modules suggests they form a constitutively active frontline defense. Furthermore, a comparison with the *E. coli* essentialome demonstrates that discrepancies in gene essentiality can often be attributed to differences in growth conditions, particularly temperature, as well as variations in genetic redundancy. In conclusion, the TNSEEK pipeline is a robust tool for exploring functional genomics across multiple strains.

**IMPORTANCE:** *Dickeya* and *Pectobacterium* are two genera of the *Pectobacteriaceae* family that contain mainly plant pathogenic bacteria. To analyze the diversity within bacteria of this family, we performed a Tn-seq analysis on six strains representing a range of ecological niches. To this aim we developed a bioinformatics pipeline, termed TNSEEK which allows the comparative analysis of results across diverse experimental conditions and multiple strains. Applied to growth in rich medium of the tested strains, it allowed the identification of an essentialome of 225 genes at the family level, few genus-specific essential genes but a large essentialome comprising 181 strain-specific genes, many of which of unknown function. Thus, TNSEEK proved its ability to analyze a large data set coming from different Tn-seq experiments and it offers an unparalleled flexibility in handling any number of strains and experiments.

## INTRODUCTION

In 2005, Samson *et al*. revised the genus *Pectobacterium*, which encompasses phytopathogenic bacteria responsible for soft rot symptoms, and established the distinct genus *Dickeya* (1). Subsequently, epidemiological and taxonomic investigations have led to the identification of numerous novel species within both genera. Currently, 23 recognized species of *Pectobacterium* and 13 species of *Dickeya* have been characterized (2, 3). Soft rot *Pectobacteriaceae* (SRP) exhibit a broad host range, encompassing both monocotyledonous and dicotyledonous plant species. The majority of these bacteria have been isolated from soft rot symptoms in a diverse array of infected plants. These plants frequently include species of significant agronomic importance, such as potato, rice, maize, cabbage, and ornamentals. These bacteria are capable of inducing disease during both pre-harvest production and post-harvest storage. Given the frequent monitoring of these crops during growth and storage, the isolation of plant-infecting strains is regularly achieved. However, certain SRP strains can also be found in alternative environments such as weeds or water (4) and some species have been mostly isolated from freshwater sources and have not been associated with plant hosts. Examples include *Dickeya aquatica*, *D. lacustris*, *D. undicola*, and *P. aquaticum* (5). Whole-genome sequencing reveals genome size variability both intragenerically and intraspecifically. Specifically, within the *Dickeya* genus, average genome sizes range from 4.02 Mb for *D. poaceiphila* to 5.35 Mb for *D. dadantii*. Furthermore, intraspecific variation in genome size is evident, with *D. zeae* exhibiting a range from 4.56 Mb to 4.93 Mb and *D. dadantii* ranging from 4.66 Mb to 5.35 Mb (3).

Genomics also allows development of high-throughput technologies, such as transposon sequencing (Tn-seq) that have facilitated the identification of numerous essential genes within a given bacterial strain (6). However, interpretations based on the analysis of a single, typically laboratory-adapted strain should be approached with caution. To mitigate this limitation, Tn-seq analysis of multiple strains is imperative. For instance, Ghomi *et al*. (7) analyzed 14 strains to define a core set of 201 essential genes for the *Enterobacteriaceae* family. Their analysis of individual strains revealed between 300 and 440 essential genes in each respective strain. Similarly, an investigation involving 17 strains of *Streptococcus pneumoniae* identified 206 core essential genes, 186 genes present in all strains but essential in only a subset, and 128 essential genes located within the accessory genome (8). These findings underscore the strain-dependent nature of gene essentiality and emphasize the necessity of analyzing and comparing data from a substantial number of strains to achieve a robust understanding of essentiality at the species, genus, and family levels.

To enable such a comparative analysis of extensive transposon sequencing (Tn-seq) datasets generated from multiple bacterial strains, we developed a fully automated bioinformatics pipeline, termed TNSEEK. This pipeline encompasses the entire data processing workflow, from raw reads to the prediction of essential genes, and permits the comparative analysis of results across diverse experimental conditions and multiple strains. TNSEEK was utilized to analyze Tn-seq data obtained from six Soft Rot *Pectobacteriaceae* (SRP) strains, comprising three *Dickeya* species (*D. dadantii*, *D. parazeae*, and *D. fangzhongdai*) and three *Pectobacterium* species (*P. atrosepticum*, *P. versatile*, and *P. aquaticum*), representing a range of ecological niches. The application of the TNSEEK pipeline facilitated the determination of the core essential genome of SRP and allowed for the differentiation of functions associated with core and strain-specific essential genes.

## MATERIAL AND METHODS

### Bacterial strains and growth conditions

All bacterial strains and oligonucleotides employed in this study are detailed in Tables S1 and S2. *Escherichia coli* cells were cultured at 37°C in Luria-Bertani (LB) medium. *Dickeya* and *Pectobacterium* cells were cultured at 25°C in 50% Tryptic Soy Broth (TSB; 20 g/L) medium (Formedium Ltd, Norfolk, UK). When required, media were solidified by the addition of 1.5 g/L agar. Antibiotics were supplemented at the following concentrations: 100 μg/mL for ampicillin (Amp), 25 μg/mL for kanamycin (Km), and 25 μg/mL for chloramphenicol (Cm).

### Construction of the transposon library

For each bacterial strain, 10 optical density units at 600 nm (OD_600_.U) of an overnight culture were combined with 10 OD_600_.U of an overnight culture of *E. coli* MFDpir/pSamEC. The pSamEC vector is a suicide mobilizable plasmid containing a kanamycin resistance gene (Km^R^) flanked by mariner inverted repeat sequences with an *MmeI* restriction site and the *himar1*-C9 transposase gene under the control of the *lac* promoter (9). This transposon specifically inserts at TA dinucleotide sites within the genome. This mixture was centrifuged at 5,000 rpm for 10 minutes at ambient temperature. The resulting bacterial pellet was resuspended in 100 µL of TSB 50% and dropped onto a TSB 50% agar plate supplemented with 57 mg/L diaminopimelate. Following incubation overnight at 30°C, bacteria were collected and resuspended in 1 mL of TSB 50%. A sample was then diluted and plated onto a TSB 50% agar plate containing kanamycin to assess mutagenesis efficiency. The remaining bacterial suspension was spread onto TSB 50% agar plates with kanamycin and incubated for 48 hours at 30°C. Bacteria were harvested from the plates by adding TSB 50%. The cultures were adjusted to an OD_600nm_ of 60, mixed with an equal volume of 80% glycerol, aliquoted, and stored at -80°C.

### DNA preparation for Tn-seq high-throughput sequencing

Genomic DNA was extracted from approximately 90 mg of bacterial cells for the control strain libraries, and from cultures grown for at least seven generations in 50% TSB at 25 °C for the mutant libraries using the Wizard Genomic DNA Purification kit (Promega), following the manufacturer’s protocol. Subsequent steps followed precisely the methodology described by Baffert *et al* (10).

### Sequencing and genome assembly

High-quality, complete genome sequences were generated using a hybrid sequencing approach, combining Oxford Nanopore Technologies (ONT) long-read and Illumina short-read sequencing. *P. versatile* strain A73-S18-O15 and *Dickeya parazeae* strain A586-S18-A17 were grown in 2 mL of liquid LB medium at 28°C. The bacterial cells were then harvested by centrifugation and genomic DNA was extracted using the method described by Sambrook and Russell (2006). Illumina sequencing was performed at the Next Generation Sequencing Core Facility of the Institute for Integrative Biology of the Cell (I2BC). Nextera DNA libraries were prepared from 50 ng of high-quality genomic DNA. Paired-end sequencing (2 × 75 bp) was conducted on an Illumina NextSeq 500/550 instrument using a High Output 150-cycle kit, resulting in approximately 60-fold coverage for strain A586-S18-A17 and 180-fold coverage for strain A73-S18-O15. Long-read sequencing was conducted at the Genotoul platform (https://www.genotoul.fr) employing Oxford Nanopore GridION technology over a 72-hour run. This yielded 447,101 reads for strain A586-S18-A17 and 335,205 reads for strain A73-S18-O15, with average read lengths of 4,473 bp and 5,173 bp, respectively. The resulting sequencing depths were 417-fold for A586-S18-A17 and 361-fold for A73-S18-O15. Hybrid genome assembly, utilizing reads from both long-read and short-read sequencing strategies, was performed using Unicycler v0.4.9 with default parameters (11). The assembled genomes were deposited and annotated on the National Center for Biotechnology Information (NCBI) platform and accession numbers for each genome are indicated in Table S1 (https://www.ncbi.nlm.nih.gov/assembly).

An orthologous gene matrix was computed for the 6 analyzed genomes using the Orthologous Matrix (OMA) standalone package (https://omabrowser.org/standalone/) (12).

Functional annotation of essential genes was performed using eggNOG-mapper v2 (13) against the eggNOG v5.0 database (14). The resulting OMA orthogroups and functional annotations are presented in Table S3.

### Description of TNSEEK

To enhance the analysis of a substantial volume of sequencing experiments, we have developed an entirely automated bioinformatics pipeline, TNSEEK, which encompasses the entire processing chain—from raw reads to the prediction of essential and conditionally essential genes. TNSEEK relies primarily on statistical approaches provided by the TRANSIT software suite (15), and offers additional features to facilitate comparison of results across multiple conditions and strains. An overview is provided in Figure 1, and its key steps are outlined below:

**Figure 1:**
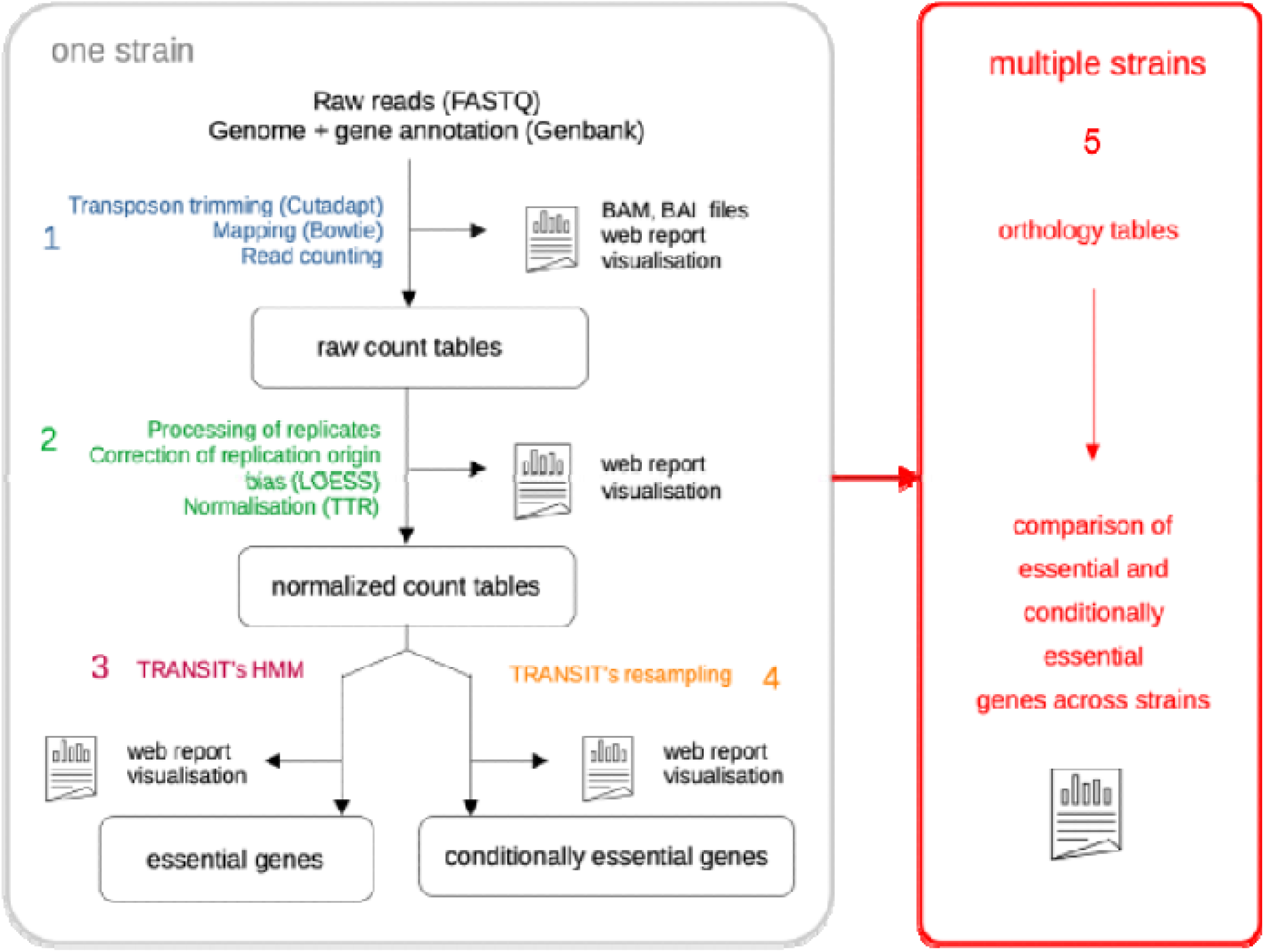
Overview of the TNSEEK pipeline. Illustration of the main steps of the pipeline, beginning with the raw sequencing reads and culminating in the identification of essential and conditionally essential genes. The pipeline initially processes data from each strain independently, followed by a comparative analysis of results across strains at the orthology family level.

Step 1: Read mapping and construction of count tables. Initially, reads were filtered to select only those containing the transposon sequence ACAGGTTGGATGATAAGTCCCCGGTCT at the 5’ end. These sequences were then trimmed from reads using Cutadapt (16), and the appropriately sized reads were mapped to the reference genome with Bowtie2 (end-to-end alignment) (17). Specific parameters are applied during this stage: the minimal overlap between a read and the transposon is set to 7 nucleotides, and a maximal error rate of 0.05 is permitted when searching for the transposon sequence. Reads are discarded if they are shorter than 16 or longer than 17 nucleotides. To build the raw count tables, only reads that align uniquely to the genome and whose 3’ end corresponds to a “TA” dinucleotide are retained; all other reads are discarded

Step 2: Normalization and quality control. We began by applying LOESS normalization (local weighted regression) to correct for replication origin bias. We then eliminated non-biological variability between tables using TTR normalization (Trimmed Total Reads). This step resulted in one normalized count table per strain and condition, along with several quality control metrics, taken from TRANSIT: sum, mean, skewness, and kurtosis of the read counts in the dataset, the density (i.e., the fraction of sites with insertions), as well as QQ (Quantile-quantile) and QC (Quality control) plots. The pipeline also performed pairwise comparisons of replicates: For each strain and each condition, we computed the Pearson correlation coefficient between TA-sites for each pair of replicates

Step 3: Detailed characterization of essential genes. Based on the read counting,_genes were classified into four categories: essential (ES), growth defect (GD), growth advantage (GA) and non-essential (NE). For that, we used the Hidden Markov Model (HMM) approach introduced in (15), that models the genome as a sequence of four hidden states calculating the likelihood of observed insertion counts for each gene.

Step 4: Characterization of conditionally essential genes: TNSEEK can also identify conditionally essential genes. TRANSIT’s RESAMPLING method is used to compare experimental conditions with the control conditions. TNSEEK features an intuitive user interface and facilitates the interpretation of results by generating a comprehensive report accessible via any standard web browser. This report enables users to systematically navigate through the analysis process, accompanied by a diverse array of figures and graphical representations. For a holistic understanding, all findings are synthesized into global tables at the strain level. This step is not used in the present study,

Step 5: Tn-seq result comparative analysis: TNSEEK provides additional insights to compare essential and conditionally essential genes across species, utilizing Orthologous Matrix Analysis (OMA) orthogroup links between strains to enhance comparative analysis. This allows to differentiate between a TNSEEK-predicted conserved, core essential genes shared between all strains and variable combinations of strain-specific essential genes, which is critical for understanding the diversity and adaptability of bacterial species.

Step 6: TNSEEK visualisation. TNSEEK also incorporates a visualization tool for the detailed examination of individual genes and comparison between orthologous genes (Fig. 2).

**Figure 2:**
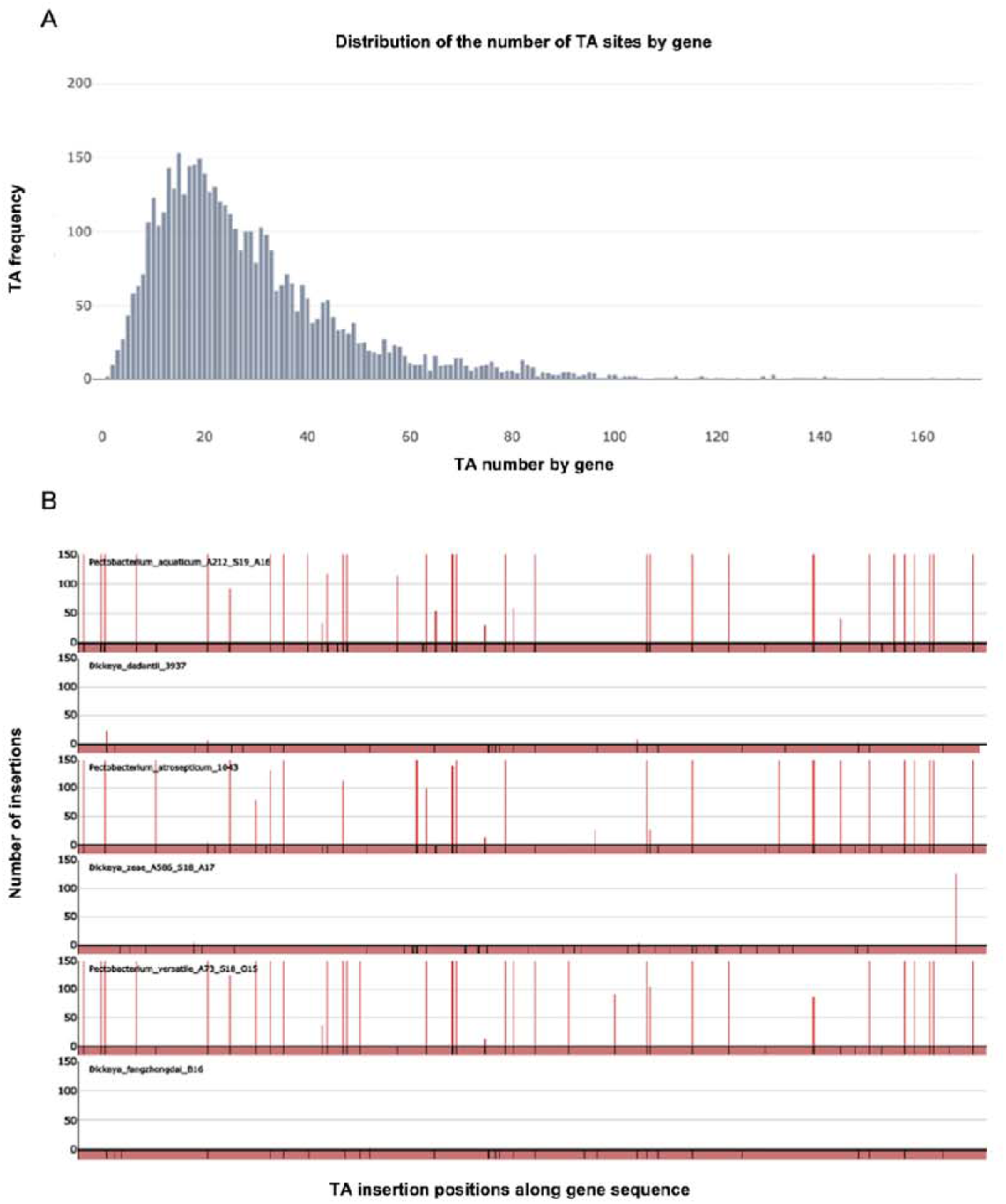
Examples of graphical outputs generated by the visualization tool of the TNSEEK pipeline. A: Distribution of the number of TA sites per gene in *D. dadantii 3937*, B: insertions visualization for an essential genus specific gene (orthogroup OMA03793). The figure clearly shows numerous insertions for the *Pectobacterium* genus (non-essential gene) and an absence of insertion for the *Dickeya* genus (essential gene). Black lines represent TA insertion sites and red lines number of insertions along the gene.

TNSEEK comes with a user-friendly interface, and facilitates result interpretation by generating a comprehensive report that can be conveniently accessed through any web browser. This report enables users to navigate through the analysis process step by step, accompanied by a diverse range of figures and graphs. For a holistic understanding, all results are summarized in global tables at the strain level, providing a comprehensive overview. Additionally, TNSEEK offers additional insights into essential genes and conditionally essential genes across species with orthology links between strains, thus facilitating comparative analysis. Lastly, TNSEEK includes a visualization tool dedicated to investigating individual genes. The computational core of TNSEEK is implemented in Python and R. The graphical user interface, specifically the web application designed for data exploration and visualization, is developed using R Shiny and SQLite. The report generation is facilitated by R Markdown. The complete pipeline is readily installable and executable as a Docker image. Source code and detailed documentation are available at https://gitlab.cristal.univ-lille.fr/bonsai/tnseek.

### Analysis of essential genes

To validate the essential gene predictions generated by the TNSEEK pipeline, several confirmatory criteria were established. Firstly, gene duplications within each strain’s genome were identified by conducting all-against-all nucleotide sequence similarity searches using BLAST 2.13.0+ (E-value threshold 10^-5^), and any duplicated genes were subsequently excluded from the essential gene list and annotated ES DU standing for “duplicated” (Table S3). Secondly, given the potential polar effects of mariner transposon insertions, the operon structure of TNSEEK-predicted essential genes was analyzed as described in the result section. UpSet plots were generated using the Python implementation of the UpSet library (Fig. 3A and 3B) (18). Intact phage regions within the six genomes were identified by PHASTEST (https://phastest.ca/).

**Figure 3:**
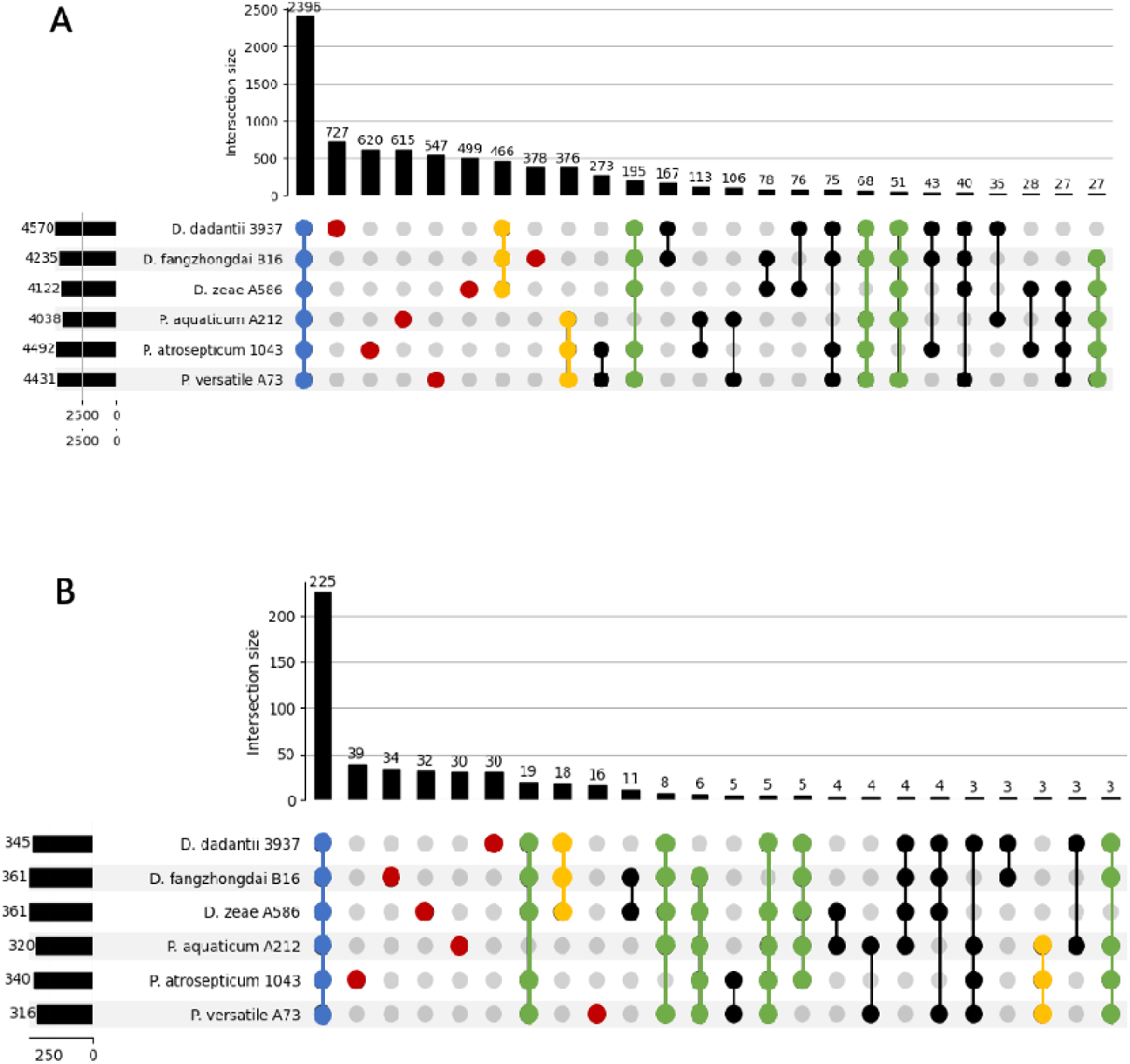
Shared genes and shared ES genes between the six studied strains. A: shared genes between the strains. B: shared ES genes between the strains. Blue: core genes shared by the 6 strains, red: strain specific genes, yellow: genus specific genes, green: genes shared by 5 strains out of 6. black: other combinations. The number of genes is indicated on the top of each bar. The graph depicted on the left represents the total number of genes (A) or ES genes (B) per strain. Only the combinations with a gene number of more than 25 are shown for (A) and of more than 2 are shown for (B).

## RESULTS

### Genomic diversity and core Genome of the six selected *Pectobacteriaceae* strains

In this study, we compared the essential genes of six SRP strains, selected to represent a variety of ecological niches. The chosen species include three from the *Dickeya* genus (*D. dadantii*, *D. parazeae*, and *D. fangzhongdai*) and three from the *Pectobacterium* genus (*P. atrosepticum*, *P. versatile*, and *P. aquaticum*). This selection encompasses a broad ecological range: *P. versatile* is isolated from both plants and surface water (2, 19), whereas *P. atrosepticum* is mainly restricted to *Solanaceae* (2), and *P. aquaticum* was found exclusively in waterways (19, 20). Similarly, within the *Dickeya* genus, *D. fangzhongdai* occupies diverse habitats, including plants and aquatic environments (21–23), *D. parazeae* is frequently found in water (19, 24) and infects herbaceous plants like maize and rice (3) and *D. dadantii* is primarily associated with dicotyledonous plants (3).

<u>T</u>wo of these strains, *P. atrosepticum* SCRI1043 and *D. dadantii* 3937, are well-established model organisms whose genomes were among the first sequenced for their respective genera (25, 26). The complete genomes of *P. aquaticum* A212-S19-A16 and *D. fangzhongdai* B16 have also been previously published (20, 23). For this study, we sequenced and assembled the complete genomes of the remaining two strains: *P. versatile* A73-S18-O15 and *D. parazeae* A586-S18-A17.

A single replicon was observed for all strains analyzed and no plasmid was detected. The genome sizes range from 4.46 Mb (*P. aquaticum*) to 5.06 Mb (*P. versatile*). The number of coding sequences (CDS) varies from 4,038 (*P. aquaticum*) to 4,570 (*D. dadantii*). The number of pseudogenes ranges from 33 in *D. fangzhondai* to 128 in *P. aquaticum*, while the number of CRISPR arrays varies from one (*P. aquaticum*) to five (*D. fangzhondai*).

While the six strains share a core genome of 2,396 genes, a substantial portion of each genome is unique (Fig 3A). The number of strain-specific genes ranges from 378 in *D. fangzhongdai* to 727 in *D. dadantii*, underscoring a high degree of genomic individuality. The analysis also reinforces the genomic divergence between the two genera, identifying 466 genes exclusive to the *Dickeya* strains and 376 genes exclusive to the *Pectobacterium* strains. As a rule, the number of genes shared exclusively in pairs between two species was higher for strains belonging to the same genus (from 273 to 76) than between species belonging to different genera (from 35 to 11). This genomic variability prompted us to investigate its functional consequences by defining and comparing the essential gene set for each strain using TNSEEK.

### Generation and validation of mariner-based transposon libraries

For the transposon sequencing (Tn-seq), a Himar9 mariner transposon derivative was utilized. A mutant pool of approximately 1,000,000 colonies was generated for each of the six SRP strains by introducing the plasposon pSam_Ec (9) via conjugation from *E. coli*. DNA libraries were prepared in two technical replicates from this pool and analyzed using high-throughput sequencing with the TNSEEK pipeline. The sequencing yielded high-quality data, with initial read counts ranging from 9.1 to 14.7 million per sample and final mapped reads between 8.6 and 13.8 million after filtering. Mapping efficiencies were consistently high, between 84.42% and 96.08%, and the mean read depth per insertion site was uniform across samples (122.2-129.4 reads), providing sufficient coverage for statistical analysis. High Pearson correlation coefficients (0.96-0.99) between replicates confirmed the experimental reproducibility. The number of unique transposon insertion sites varied between the genera, with *Pectobacterium* strains showing higher counts (199,301–232,639) than *Dickeya* strains (166,816–194,008), reflecting differences in genome size and saturation. The TA insertion density, ranging from 37.9% to 62.7%, was sufficient for a comprehensive determination of gene essentiality. The use of a standardized growth condition (TSB 50%) allowed for the direct comparison of essential gene repertoires, which facilitates the identification of both conserved core essential genes and species-specific fitness determinants among these related phytopathogenic bacteria.

### Defining the core and variable essentialome of *Pectobacteriaceae*

The initial step in our functional analysis involved processing the raw Tn-seq data with the TNSEEK pipeline to identify essential genes in each of the six strains. This first-pass analysis yielded a set of 665 orthogroups containing at least one potentially essential gene (Table S3). However, a known challenge with transposon-based methods like Tn-seq is the risk of generating false positives due to polar effects (27). An insertion of the mariner transposon within a gene can not only disrupt that gene but also prevent the transcription of downstream genes located in the same operon. This can make these downstream genes appear essential, when in reality their function was never directly tested by an insertion.

To address this challenge and ensure the high quality of our dataset, we implemented a stringent and systematic curation strategy. The logic of our approach was to specifically identify and remove cases where essentiality is questionable due to a polar effect. We leveraged both known operon structures for *D. dadantii* 3937 (28) and predicted operon structures based on synteny for the other five strains. For each putative operon containing multiple genes identified as essential, we applied a conservative rule: only the gene located most distal from the promoter was retained as essential. The rationale is that an insertion in this last gene is the least likely to have a polar effect on any other gene, making its essentiality call the most reliable. However, this rule was applied with a critical exception to avoid inadvertently discarding truly essential genes. If a gene located upstream in an operon had a known essential ortholog in the well-characterized model organism *E. coli* K-12 (29) was maintained in our essential gene set. This bioinformatic cross-validation allowed us to anchor our predictions in established biological knowledge. This meticulous approach enabled us to identify 455 orthogroups whose essentiality in at least one of the six analyzed strains could not be attributed to polarity effects. Among the 210 orthogroups categorized as ES by the TNSEEK pipeline that could be prone to artifactual categorisation due to the polarity of the Mariner transposon, 97 were categorized as essential in *E. coli* K12 by Goodall et al (29). We opted to include these 97 orthogroups in our subsequent analysis. The majority of these, 76 orthogroups, were categorized as ES across all six analyzed strains. Through this rigorous, multi-step curation process, we successfully filtered out ambiguous cases. This refined our initial list to a final, high-confidence dataset of 552 orthogroups containing essential genes, with at least one ES gene across the six analyzed strains (Table S4). This curated dataset formed the robust foundation for the subsequent comparative analysis of the core and variable essentialomes.

### The comparative analysis of the 6 strains discriminates between core, genus-specific and strain-specific essentialomes

Our comparative analysis, based on the high-confidence set of 552 essential orthogroups, highlights a distinction in the genetic requirements for survival among the SRP strains. We first identified a conserved core essentialome comprising 225 orthogroups (Figure 3B and Table S5). These genes are required for growth in all six strains under the tested conditions. The total number of essential genes per strain was also relatively consistent, varying from 316 for *P. versatile*, 320 for *P. aquaticum*, 345 for *D. dadantii*, 340 for *P. atrosepticum* and 361 for *D. fangzhongdai* or *D. parazeae*.

In addition to this core set, we identified a large strain-specific essentialome, consisting of 181 genes pointing toward a considerable degree of functional individuality. The contribution of this variable set to each strain’s essentialome differed, ranging from 16 unique essential genes in *P. versatile* to 39 in *P. atrosepticum* (Figure 3B and Tables S6, S7, S8, S9, S10, S11). This suggests that each strain may have developed specific dependencies on certain accessory genes. Only a few orthogroups exhibited genus-specific essentiality: 18 orthogroups were found to be exclusively essential for the three *Dickeya* species (Table S12). Even fewer, 3 orthogroups, were exclusively essential for the three *Pectobacterium* species (Table S13 and Figure 3B).

We then investigated the genetic basis for this strain-specific essentiality. In a high proportion of cases (87.6%), a gene was essential in a single strain because its orthologs were absent from the genomes of the other five strains (Table 1). This observation links the functional essentiality to the genomic diversity described previously, indicating that the variable essentialome is primarily composed of the unique gene repertoire of each strain. Furthermore, our analysis suggests that gene essentiality can be viewed as a continuum of fitness contribution. This was apparent for genes essential in a majority of strains (four or five). In these instances, their orthologs in the remaining strains were predominantly classified as causing a growth defect (GD) (Table 1). This indicates that while the absence of these genes is not strictly lethal in all genetic backgrounds, it likely imposes a significant fitness cost.

**Table 1:** Status of the remaining genes (%) not categorized ES by TNSEEK when at least one gene of the OMA orthogroup is essentiel.

| Nb of ES essential genes within the OMA orthogroup | Status of the non ES genes* (% total) |  |  |  |
| --- | --- | --- | --- | --- |
|  | Growth defect | non essential | Absent | Duplicated |
| 6 | 0.0 | 0.0 | 0.0 | 0.0 |
| 5 | 76.1 | 13.0 | 8.7 | 2.2 |
| 4 | 87.5 | 12.5 | 0.0 | 0.0 |
| 3 | 47.2 | 37.0 | 13.9 | 1.8 |
| 2 | 39.7 | 39.7 | 19.9 | 0.6 |
| 1 | 10.4 | 26.7 | 87.6 | 0.4 |

\*The Growth advantage category is not represented in the table as none of the non-ES gene fall into this category

In summary, the essentialome of these *Pectobacteriaceae* strains is characterized by two components: a stable, conserved core reflecting their shared biology, and a larger, variable component derived from the accessory genome, which appears to shape the specific survival dependencies of each strain.

### Distinct functional signatures of the core and strain-specific essentialomes

To investigate the biological roles of the core and variable essentialomes, we performed a functional analysis using the COG (Clusters of Orthologous Groups) database. The results revealed clear differences in the functional profiles of these two gene sets (Figure 4A).

**Figure 4:**
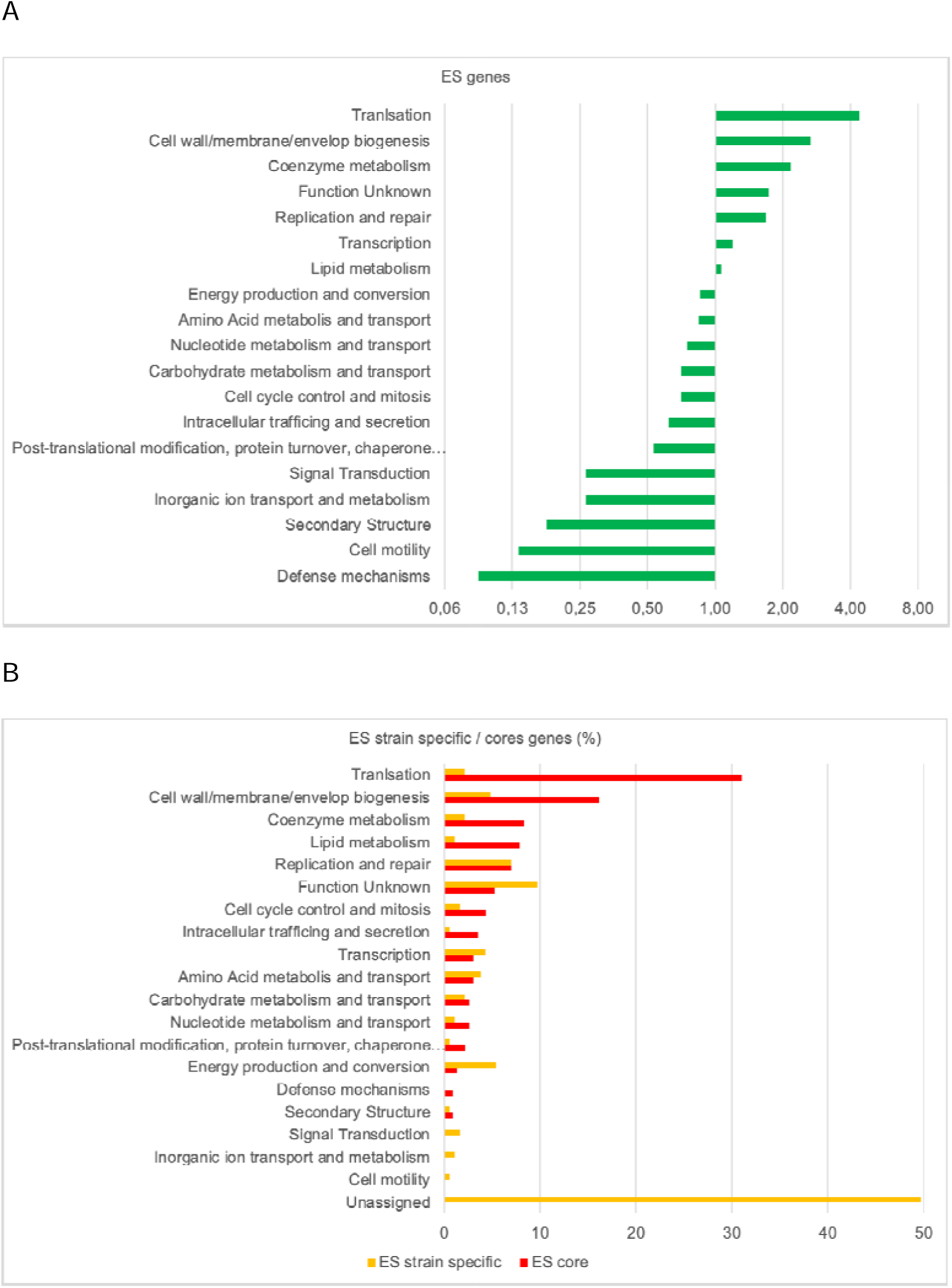
Distribution of COG categories. A: representation factor of all essential genes (ES) according to total genes for each COG category (Log2 scale). Factors above 1 mean over-representation, below 1 under-representation according to the total genes. B: percentage of COG categories for the 181 strain specific ES genes (yellow bars) or the 225 core ES genes (red bars). Among the strain specific ES genes nearly 50% are not assigned to any COG category.

First, we analyzed the functional distribution of all 552 orthogroups containing at least one essential gene. Of these, 458 could be assigned to a COG category, while 94 remained unclassified. As expected, categories related to core cellular processes were significantly enriched. Specifically, the most overrepresented functions included “Translation, ribosomal structure and biogenesis” and “Cell wall/membrane/envelope biogenesis”. In contrast, functions related to environmental interaction and adaptation, such as “Defense mechanisms” and “Cell motility”, were the most underrepresented. A contrast was observed when comparing the 225 core essential genes with the 181 strain-specific essential genes (Figure 4B). The profile of the variable essentialome was different. A high proportion (50%) of the 181 strain-specific essential genes could not be assigned to any COG category, suggesting a gap in the functional annotation of these genes. For the genes that could be classified, the most frequent designation was “Function unknown”, which accounted for over 10% of the set. This functional divide indicates that while the machinery for cell maintenance appears conserved, a portion of each strain’s specific dependencies may be governed by a diverse and less-characterized collection of genes.

### Comparative analysis with *E. coli* K-12 reveals the influence of growth conditions and functional redundancy

The *Pectobacteriaceae* family is part of the *Enterobacterales* order (30). To better understand the factors governing gene essentiality in *Pectobacteriaceae*, we compared the essentialome of the six SRP strains with that of the well-characterized *Enterobacterales* model, *E. coli* K-12. This comparison allowed us to identify genes with differential essentiality and to explore the biological reasons for these differences.

Our analysis identified a set of 24 genes that are considered essential for *E. coli* K-12 but were found to be non-essential in all six of the SRP strains we investigated (Table S14). A closer examination of these genes suggests that these discrepancies are not due to fundamental biological divergence but rather highlights the principle of conditional essentiality, where a gene’s requirement is dependent on the specific environmental context. A clear illustration of this is the *cydABCDX* gene cluster. In *E. coli*, these genes encode the cytochrome bd-I quinol oxidase, a terminal oxidase in the electron transport chain. Cytochrome bd-I is assembled from three core proteins: CydA (orthogroup OMA0072), CydB (OMA01799), and CydX (QR68_06300). In addition, two accessory proteins, CydC (OMA00690) and CydD (OMA00639), function as cysteine and glutathione exporters. These transporters are essential for cytochrome maturation, as they facilitate the incorporation of heme (31). While this complex is not required for growth at moderate temperatures, it becomes critical for survival and cell division at 37°C, the standard culture temperature for *E. coli* (32). Consequently, it is classified as essential under these conditions. However, our SRP strains were cultured at a lower temperature of 25°C. At this temperature, alternative respiratory pathways are sufficient in *E. coli*, and the *cyd* genes are not essential. The non-essential status of these genes in our dataset is therefore consistent with the known temperature-dependent physiology of *Enterobacterales*. A similar temperature-dependent essentiality is observed for the *rpoE* gene. An *E. coli rpoE* mutant is unable to grow above 30°C, rendering *rpoE* essential at 37°C (33). Importantly, *Dickeya* and *Pectobacterium rpoE* mutants exhibit no growth defects at 25°C, the temperature used in our experiments.

Furthermore, it is plausible that genes causing non-lethal growth defects in *E. coli* at 37°C may be classified as essential in our study due to the different growth conditions. For instance, genes such as *nlpD* (OMA02130), *priA* (OMA00249), and *secF* (OMA03091) are classified as non-essential in Goodall’s *E. coli* dataset (29). However, mutations in these genes are known to impair growth, with the *secF* mutant, for example, displaying exacerbated defects at lower temperatures (34–36). It is therefore possible that such growth defects are more pronounced in *Dickeya* and *Pectobacterium* when cultured at 25°C, leading to the depletion of these mutants from the population and thus an essential phenotype under our experimental conditions.

Conversely, our analysis identified 14 orthogroups essential in all six SRP strains but non-essential in *E. coli* K-12 (Table S15). The presence of paralogs and the existence of non-orthologous functional analogs in *E. coli* account for these discrepancies. For example the six SRP strains possess only a single copy of the *gpmA* essential gene (OMA03987), which encodes a phosphoglyceromutase. In contrast, the *E. coli* genome contains in addition to *gpm*A the paralog *gpmM*, that performs the same function. Due to this duplication, *E. coli* can tolerate the loss of *gpmA*, as the function is maintained by *gpmM*. The *gpmA* gene is therefore classified as non-essential in this genomic context. Both genes must be disrupted simultaneously in *E. coli* to inhibit growth (37). The second mechanism is the use of non-orthologous functional analogs, where the two organisms utilize different proteins to perform the same essential task. This is the case for the DNA helicase loader function, a critical step in the initiation of DNA replication. In *E. coli*, this essential function is encoded by the *dnaC* gene (38). The SRP strains rely on a different helicase, belonging to orthogroup OMA05632, which is essential in all six strains (39). Thus, although the function is essential in both groups, it is performed by evolutionarily distinct genes.

#### Genus-specific patterns of gene essentiality suggest a few distinct metabolic capabilities between *Dickeya* and *Pectobacterium* genera

Eighteen orthogroups contain genes that are essential in the three *Dickeya* species but not in the three *Pectobacterium* species (Table S12). A clear example is the D-alanine-D-alanine ligase (Ddl), which catalyzes a crucial step in peptidoglycan synthesis. *Dickeya* species possess only the *ddlA* gene that codes for this activity (orthogroup OMA02306), while *Pectobacterium* species have only *ddlB* (OMA03351). Consequently, the deletion of *ddlA* is lethal in *Dickeya*, and the deletion of *ddlB* is lethal in *Pectobacterium*. In contrast, *E. coli* possesses both *ddlA* and *ddlB*, providing functional redundancy that allows it to survive the loss of either gene individually (40). Another case involves the *nrdA* (OMA00091) and *nrdB* (OMA01483) genes, which encode the two subunits of the class Ia ribonucleotide reductase. These genes are essential in both *E. coli* and *Dickeya*, but surprisingly, they are not essential in *Pectobacterium* (Table S12) (Fig. 5A). While no duplication of *nrdA/B* was found in *Pectobacterium*, they do possess an alternative ribonucleotide reductase system, encoded by *nrdE* and *nrdF*. This alternative enzyme, which is absent in *Dickeya*, is known to function in *E. coli* under specific conditions, such as iron-depleted, manganese-rich conditions (41). Further investigation is needed to confirm if the *nrdE/F* system provides a functional substitute for *nrdA/B* in *Pectobacterium* under the growth conditions used in this study.

**Figure 5:**
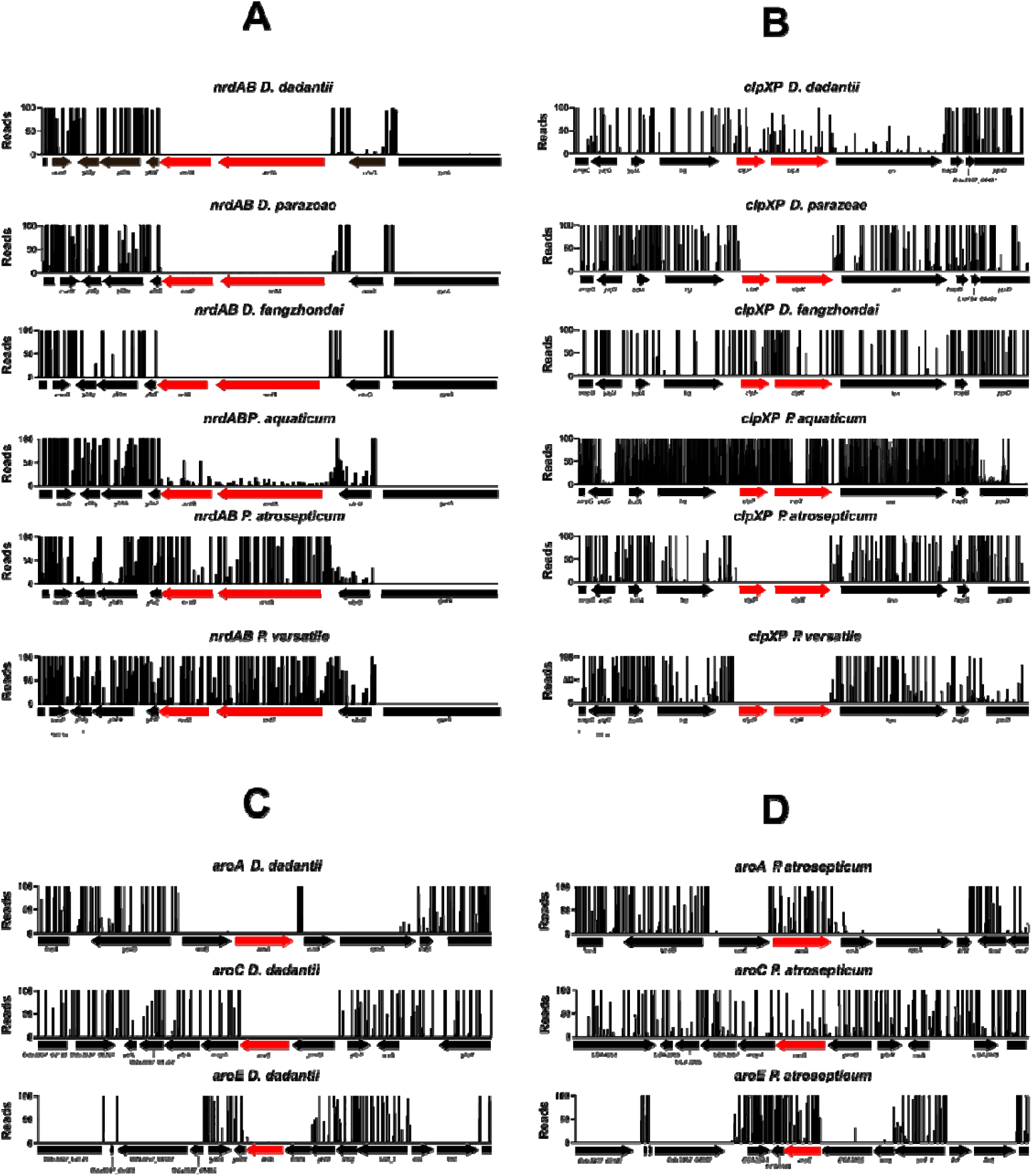
Tn-seq insertion profiles at selected loci in various *Dickeya* and *Pectobacterium* species. Each panel shows normalized Tn-seq read counts (y-axis, “Reads”) mapped along the genomic region encompassing the indicated locus. Black peaks represent mapped insertion reads; absence of peaks indicates a lack of Tn insertions due to essentiality under the tested growth conditions. Red arrows highlight the gene(s) assessed for essentiality in each panel: (A) *nrdAB* cluster; (B) *clpXP* operon; (C) *aroA*, *aroC*, and *aroE* in *D. dadantii*; (D) *aroA*, *aroC*, and *aroE* in *P. atrosepticum*. Gene organization is depicted with black arrows and gene names below each region.

**Figure 6:**
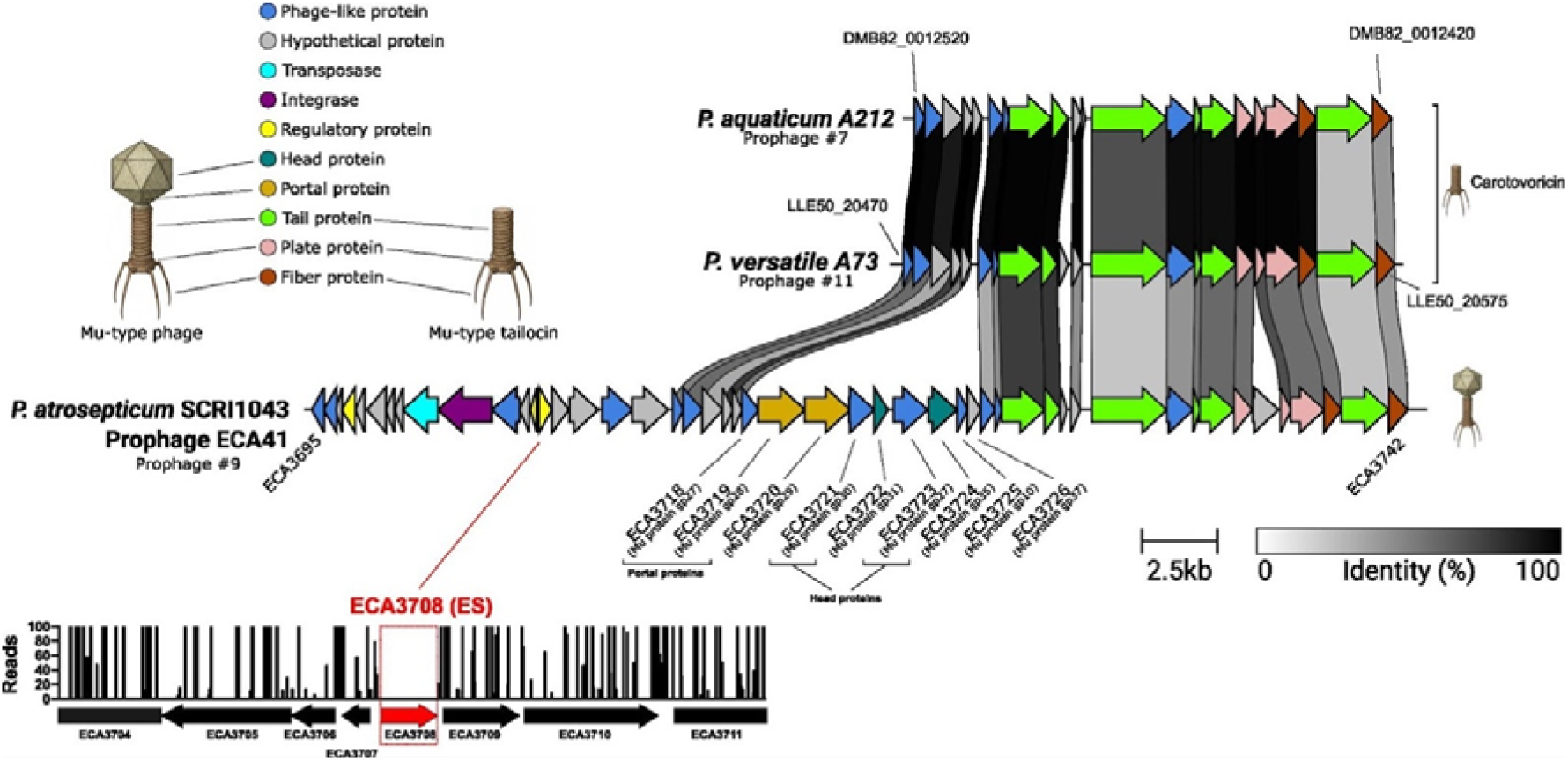
Comparative genomic analysis of a functional Mu-like prophage and two defective tailocin-encoding prophages in *Pectobacterium* species. Genomic alignment, generated using clinker, of prophage regions from three *Pectobacterium* strains. The bottom line displays the Mu-like prophage #9 (ECA41) from *P. atrosepticum* SCRI1043. This prophage contains the essential (ES) regulatory gene *ECA3708*, highlighted in red, which is required to maintain lysogeny. Genes encoding structural proteins, such as portal and head components, are also annotated. In contrast, the middle and top lines show prophage #11 from *P. versatile* A73 and prophage #7 from *P. aquaticum* A212. These regions are defective prophages that constitute the carotovoricin operons, which encode a Mu-type tailocin, a potent antibacterial weapon. Arrows represent predicted genes, colored by their putative function according to the key. Gray links between the regions illustrate amino acid identity, with the color intensity corresponding to the percentage of identity. The bottom histogram shows the distribution of Tn-seq reads mapped across the prophage region in *P. atrosepticum*, reflecting the transposon insertion profile. The absence of insertions (i.e., no reads) in the essential gene ECA3708 (highlighted in red) indicates its requirement for bacterial viability under tested conditions, while neighboring genes with higher insertion counts are dispensable.

Conversely, a more limited set of genes was identified as essential specifically within the *Pectobacterium* genus. Only three orthogroups contain genes that are essential in the three studied *Pectobacterium* species but not in the three *Dickeya* species (Table S13). OMA04333 includes essential orthologous genes in each *Pectobacterium* species (DMB82_0011620, ECA2435, and LLE50_21950). Based on sequence similarity and domain architecture, these proteins are orthologs of RdgA, a regulator previously identified in *P. versatile* strain 71. RdgA is linked to the control of pectin lyase production and, by homology with *E. coli*, to DNA repair and stress-response pathways (42). We hypothesize that these RdgA orthologs act as stress-responsive repressors to coordinate DNA damage repair, control expression of potentially deleterious genes during rapid growth in rich media, and thereby maintain cellular homeostasis and viability. Given their essentiality in *Pectobacterium*, experimental characterization of their precise role is warranted.

Distinct essentiality patterns of shikimate pathway genes were also observed between *Dickeya* and *Pectobacterium* when grown in TSB 50%. Specifically, in *Dickeya,* the genes *aroA*, *aroC*, and *aroE* are essential, while these same genes are non-essential in *Pectobacterium* (Fig. 5C and 5D). These genes encode critical enzymes responsible for aromatic amino acid biosynthesis, culminating in the production of chorismate that is vital for synthesizing phenylalanine, tyrosine, tryptophan, and other essential metabolites. Additional observations show that *aroF*, encoding a tyrosine-sensitive DAHP synthase catalyzing the first committed step, exhibits a growth-defect phenotype in *Dickeya* but is non-essential in *Pectobacterium*, while *aroQ*, encoding a type II 3-dehydroquinate dehydratase, remains essential in both genera. Two other DAHP synthase isoenzyme genes, *aroG* and *aroH*, as well as the aromatic amino acid permease gene *aroP*, are non-essential in both. This pattern suggests that *Dickeya* more heavily depends on *de novo* aromatic amino acid biosynthesis for growth, even in nutrient-rich conditions, whereas *Pectobacterium* compensates for pathway disruption effectively, likely through superior transport and utilization of environmental amino acids and peptides. Proposed hypotheses to explain these genus-specific differences include greater abundance or affinity of aromatic amino acid transporters and enhanced peptide uptake and hydrolysis systems in *Pectobacterium*, allowing it to scavenge sufficient aromatic amino acids from the complex peptides in TSB medium. Additionally, evolutionary divergence in metabolic regulation and differences in the demand for chorismate-derived compounds may contribute to these distinct physiological strategies.

### Defense related systems reveal strain-specific essential Restriction-Modification (R-M), Toxin-Antitoxin and contact-dependent growth inhibition (CDI) systems

Defense-related systems are crucial for protecting bacterial cells from foreign genetic elements like bacteriophages and plasmids. To systematically identify the essential genes carried by these modules, we analyzed all six genomes using the Defense-Finder tool. This analysis revealed that the vast majority of defense genes are non-essential for growth in rich medium. Specifically, out of 197 identified genes belonging to anti-phage systems, 152 were categorized as non-essential. Nevertheless, by comparing the predicted defense systems (Table S16) with our catalog of essential genes, we identified a critical subset of 14 essential anti-phage genes belonging to eight distinct system types. This finding suggests that the molecular activity associated with these essential genes is constitutive under our experimental conditions, thereby imposing a selective pressure for their function even in a phage-free environment.

Several of these essential genes are part of active Restriction-Modification (R-M) systems. In these systems, a methyltransferase enzyme modifies specific DNA sequences in the host genome, protecting it from cleavage by a corresponding restriction endonuclease. This mechanism becomes essential for cell survival if the restriction enzyme is constitutively expressed. A clear example is found in *D. fangzhongdai*, where the gene QR68_02190, encoding a DNA cytosine methyltransferase, was identified as essential (Table S16). This gene is located adjacent to QR68_02195, which encodes the restriction enzyme MvaI which recognizes the sequence 5′-CCAGG-3′ and is inhibited by the methylation of the internal cytosine residue (43, 44). The essentiality of the methyltransferase (QR68_02190) implies that the MvaI restriction enzyme is actively expressed in the TSB 50% medium, making protective methylation a prerequisite for survival. A similar configuration was observed in *D. parazeae*, where the essential gene LHK94_01575 encodes a site-specific DNA methyltransferase (tableS16). This enzyme is homologous to the SmaI methyltransferase from *Serratia marcescens*, which targets the palindromic sequence 5′-CCCGGG-3′. Located in its genomic vicinity is LHK94_01580, which encodes a Type II restriction enzyme homologous to the SmaI enzyme. Therefore, the essentiality of the methyltransferase gene LHK94_01575 is required to prevent the LHK94_01580 enzyme from cleaving the chromosome. These two examples show that specific Type II R-M systems are essential in *D. fangzhongdai* and *D. parazeae*. This contrasts with many other methyltransferases identified by Defense-Finder, which were classified as non-essential in our analysis. For these non-essential systems, it is likely that their cognate restriction enzymes are either absent from the genome or not actively expressed under the tested conditions.

The analysis of defense systems revealed that several Toxin-Antitoxin (TA) systems exhibit diverse essentiality profiles across the studied strains (Table S16). These systems typically function as addiction modules to ensure their stable inheritance. The essentiality of an antitoxin gene is a strong indicator that its corresponding toxin is actively expressed, making the antitoxin’s neutralizing function critical for host viability. Defense Finder also identified several Toxin-Antitoxin (TA) systems with diverse essentiality profiles. Among these, the RosmerTA system consists of the RmrA antitoxin, a Zn-dependent peptidase, and the RmrT toxin, which induces membrane depolarization (45). While the RmrT toxin shows sequence variability, the TA system is conserved in *D. parazeae* (LHK94_20535/LHK94_20530), *P. atrosepticum* (ECA2912/ECA2211), and *P. versatile* (LLE50_14920/LLE50_14925). In these three species, the *rmrA* antitoxin gene was essential, strongly suggesting that the RmrT toxin is active under our experimental conditions. Intriguingly, RmrT toxins have previously been shown to be responsible for the essentiality of the ClpX chaperone in *S. pneumoniae* (46) and the ClpP protease in *E. coli* E101 (47). Our results reveal a compelling correlation that supports this model: the genes encoding the ClpX protease (OMA01412) are essential exclusively in the three strains (*D. parazeae*, *P. atrosepticum*, and *P. versatile*) that contain the RosmerTA system (Fig 5B).

Another essential module identified is the AbiE system, a Type IV TA system (48) present in *P. aquaticum* and *P. versatile* (Table S16). This system is encoded by a bicistronic operon that includes the toxin AbiEii, a GTP–binding nucleotidyltransferase that causes reversible growth arrest, and the antitoxin AbiEi, which acts as a transcriptional repressor by binding the *abiE* promoter. In both *P. aquaticum* and *P. versatile*, the antitoxin genes (DMB82_0015475 and LEE50_14595, respectively; AbiEi_3-orthogroup OMA07585) were found to be essential, indicating active expression of the corresponding toxin genes (DMB82_0015470 and LLE50_14590; AbiEii-orthogroup OMA06941). In contrast, *D. parazeae* possesses a homologous antitoxin gene (LHK94_08840) that is non-essential. This is explained by the absence of the cognate toxin gene of the orthogroup OMA06941 in its genome (Table S3).

A final example found by DefenseFinder involves a Type II Toxin-Antitoxin system belonging to the RelE/ParE family. In *P. aquaticum*, the gene (DMB82_0003135), which encodes an antitoxin characterized by a C-terminal XRE-HTH domain, was identified as essential (table S16). This antitoxin is associated with its cognate toxin encoded by the adjacent gene (DMB82_0003140). The essentiality of this antitoxin is conserved in related species possessing homologous TA systems, such as *P. atrosepticum* (ECA0673/ECA0674) and *D. fangzhongdai* (QR68_07805/QR68_07810). A notable exception is observed in *P. versatile*, where the antitoxin gene (LLE50_14865) is non-essential despite the presence of its cognate toxin gene (LLE50_14870), suggesting a different regulatory mechanism or lack of toxin expression under the studied conditions.

This principle of an immunity protein being essential to neutralize a cognate toxin extends beyond canonical Toxin-Antitoxin systems to those involved in intercellular competition. In the *D. dadantii* 3937 genome, three ES genes, *Dda3937_02097*, *Dda3937_01757*, and *Dda3937_04645*, unique to *D.dadantii* 3937, encode immunity proteins that protect the producing strain from self-intoxication by contact-dependent growth inhibition (CDI) systems (Table S6). *Dda3937_02097*, also known as *cdiI,* encodes an immunity protein that neutralizes the CdiA toxin delivered via a classical CDI system (49). Similarly, *Dda3937_01757* (*rhsIA*) and *Dda3937_04645* (*rhsIB*) encode immunity proteins specific to RhsA and RhsB toxins, respectively (50). The essential function of the 3 genes strongly indicates that their cognate toxins are actively expressed under the tested conditions and exert a toxic effect in the absence of their immunity partners.

### Essential genes within prophage regions suggest lysogeny and defensive roles

To assess the presence of potentially essential genes within prophage regions, we employed the PHASTEST tool (51) to analyze the six SRP genomes. PHASTEST identified a total of 13 intact prophage regions across the six genomes: one in *D. fangzhongdai*, two each in *D. dadantii* 3937, *D. parazeae* A586, *P. aquaticum* A212, and *P. atrosepticum* SCRI1043, and four in *P. versatile* A73 (Table 2 and S17). Prophages #2, 3# #7 and #11 are defective prophages encoding tailocins, bacterial phage tails active as antibacterial weapons conferring a significant competitive advantage within the plant niche (52). Prophages #8 and #9 of *P. atrosepticum* have been studied by Evans *et al.* (53) and are named ECA29 and ECA41, respectively. The number of essential genes per prophage varied from zero to four, with 11 out of the 13 prophages harboring at least one essential gene (Table 2 and S17).

**Table 2.** : **phage essential genes**

| Strain | Prophage number/<br>name | Region length<br>(kb) | Region Position | First locus tag* | Last locus tag* | Number of ES genes | Putative phage repressor |  | List of ES genes |
| --- | --- | --- | --- | --- | --- | --- | --- | --- | --- |
|  |  |  |  |  |  |  | description | locus tag |  |
| *D. dadantii* 3937 | 1 | 38 | 915372-952946 | Dda3937_01762 | Dda3937_02545 | 4 | C1 | Dda3937_01765 | Dda3937_01763<br>Dda3937_01765<br>Dda3937_02545<br>Dda3937_02546 |
|  | 2 Dickeyocin<sup>a</sup> | 28 | 2733074-2760902 | Dda3937_03777 | Dda3937_00104 | 1 | C1 | Dda3937_03777 | Dda3937_03777 |
| *D. parazeae* A586 | 3 Dickeyocin<sup>a</sup> | 25 | 1993353-2018459 | LHK94_18375 | LHK94_18525 | 1 | C1 | LHK94_18525 | LHK94_18525 |
|  | 4 | 59 | 2446385-2505189 | LHK94_20390 | LHK94_20710 | 2 | XRE-family<br>HTH<br>Transcriptional<br>regulator,<br>peptidase s24<br>domains | LHK94_20615 | LHK94_20495<br>LHK94_20530 |
| *D. fangzhongdai* B16 | 5 | 28 | 2806765-2834379 | QR68_12205 | QR68_12360 | 1 | C1 | QR68_12205 |  |
| *P. aquaticum* A212 | 6 | 55 | 662735-718114 | DMB82_000312<br>5 | DMB82_000345<br>0 | 3 | C1 | DMB82_0003205 | DMB82_0003135<br>DMB82_0003185<br>DMB82_0003205 |
|  | 7<br>carotovoricin<sup>a</sup> | 20 | 2653251-2673677 | DMB82_001242<br>0 | DMB82_001252<br>0 | 0 |  |  |  |
| *P. atrosepticum*<br>SCRI1043 | 8 ECA29<br>prophage<sup>b</sup> | 31 | 2□935□264<br>-<br>2□966□783 | ECA2598 | ECA2637 | 1 | C1 | ECA2636 | ECA2636 |
|  | 9 ECA41<br>prophage<sup>b</sup> | 36 | 4□144□591<br>-<br>4□180□799 | ECA3695 | ECA3742 | 1 | helix-turn-helix-XRE -<br>transcriptional<br>regulator | ECA3708 | ECA3708 |
| *P. versatile* A73 | 10 | 36 | 217090-252597 | LLE50_12330 | LLE50_12560 | 2 | C1 | LLE50_12345 | LLE50_12345<br>LLE50_12555 |
|  | 11<br>carotovoricin<sup>a</sup> | 22 | 2022299-2043834 | LLE50_20470 | LLE50_20575 | 0 |  |  |  |
|  | 12 | 47 | 3215138-3261670 | LLE50_03075 | LLE50_03375 | 1 | C1 | LLE50_03110 | LLE50_03110 |
|  | 13 | 37 | 3595501-3632713 | LLE50_04915 | LLE50_05140 | 2 | helix-turn-helix-XRE -<br>transcriptional<br>regulator | LLE50_05055 | LLE50_05050<br>LLE50_05055 |

\*for *D. dadantii* 3937, the locus tags are not ordered
<sup>a</sup> Evans et al, 2010 , <sup>b</sup> Borowicz et al 2025

The most frequently identified essential gene function was prophage repression. Specifically, essential CI-type repressors were found in eight prophage regions. CI repressors maintain the prophage in its lysogenic state by preventing the expression of genes required for the lytic cycle. Loss of CI function would typically trigger prophage excision and cell death via lysis. For instance, the essential classification of *D. dadantii* 3937 gene *Dda3937_01765*, which encodes a CI repressor, directly reflects its role in preventing the lytic cascade. Furthermore, three other prophage regions harbor an essential gene encoding a helix-turn-helix (HTH) XRE transcriptional regulator. XRE-type regulators are a broad family of bacterial proteins known to modulate gene expression via HTH DNA-binding domains and are often involved in various processes, including phage–host interactions. These XRE proteins likely serve a function analogous to the CI repressor (Table 2) (54, 55). Collectively, these findings suggest that 11 of the 13 identified prophage regions likely encode functional lysogenic phages and dickeyocin. Beyond the repressors, other essential genes were identified in the prophage regions (Table 2). For instance, the *Dda3937_01763* gene (prophage #1) encodes an ASCH domain-containing protein with endoribonuclease activity, potentially acting to prevent virion production. Additionally, a functional Toxin-Antitoxin pair was identified in prophage #10 of *P. versatile* (Table S16), where the encoded antitoxin (*LLE50_12555*) is essential.

## DISCUSSION

Tn-seq is a powerful tool to study bacterial strain fitness in various environments. However, to disentangle strain-specific essential fitness traits from those affecting more broadly the studied species, genus, or family it is imperative to study and compare several strains. For this purpose, we created the TNSEEK pipeline that allowed a straightforward and easy comparison of all the Tn-seq data. We used TNSEEK to compare the results of Tn-seq experiments performed on TSB 50% for six strains belonging to two genera within the *Pectobacteriaceae* family.

The core essentialome of the *Pectobacteriaceae*, that regroups 225 essential genes conserved in the six studied bacteria, has almost the same size as the ones defined for the *Enterobacteriaceae* family (201 genes/13 strains from 5 genera) (7). Interestingly, the core essentialome defined for the single species *Streptococcus pneumoniae* (206 genes/36 strains studied) is also of a similar size (8). This suggests a lack of connection between essentiality and phylogeny, as was previously observed in another study (47). Furthermore, our analysis indicates that analyzing just six strains is enough to address this essential core at the family level as already observed at the species level for *P. aeruginosa* (56).

Our work also confirmed that, following Tn-seq analysis, essentiality can be viewed as a fitness continuum. Indeed, the TNSEEK pipeline revealed that most genes that were essential in four to five of the analyzed strains were categorized as growth defects (GD) in the other tested strains. This is due to the fact that, in a Tn-seq screen, each mutant competes with the population of bacterial cells that are functional for the same character. Therefore, most mutations that lead to a loss of fitness will either decrease in frequency or be eliminated from the population, resulting in GD or ES categorization, respectively.

Comparing our Tn-seq analysis with one performed on *E. coli* K12 provided potential explanations for some of the differences between the two studies. Firstly, while a previous study suggested that a shift in essentiality is not associated with a shift in expression level (47), we found that the choice of growth conditions, particularly temperature, can significantly influence the determination of the essential gene set and account for some of the observed results. Secondly, duplication events occurring in one strain but not the other could also influence the outcome of Tn-seq experiments. In-depth analysis also supports this latter explanation for some of the differences in essentiality observed between the six strains studied. Additionally, alternative enzymes that fulfil the same essential function, or differential gene regulation between strains, were also identified as potential explanations for the differences between the two studies.

Our comparative analysis also identified a large set of 181 species-specific essential genes, ranging from 39 in the case of *P. atrosepticum* to 16 in the case of *P. versatile*. Most of these species-specific ES genes are absent from the genomes of the other five analyzed species, which highlights the importance of variable genomes for essentiality. As only one strain was studied per species, it was not possible to determine whether the observed specificity is linked to the analyzed species or the specific strain studied. Future Tn-seq analyses involving a set of strains from the same species could distinguish between ES-specific genes that are conserved at the species level and those that are strain-specific. Interestingly, COG analysis did not assign most of these ES-specific genes to any known function. This contrasts with core essential genes, which are mostly associated with the maintenance of bacterial cell functions. The essentiality of a large number of genes with unknown functions highlights the importance of conducting functional studies in future.

Our focused analysis of the defensome indicated that a few of the ES gene functions were related to various defense mechanisms including restriction/modification systems, toxin/antitoxin mechanisms and CrispR proteins for example. The essentiality of these defense mechanisms in TSB 50%, in the absence of known stressor or bacterial competition, suggests that they form a constitutive front line that is deployed in the absence of immediate danger, and could be further reinforced under stress or attack by inducible defense mechanisms, since most of the defensome proteins remain non-essential. Notably, the mechanisms responsible for this constitutive frontline defense vary greatly among the tested strains, highlighting its high evolvability. These species-specific patterns suggest that distinct regulatory architectures, ecological pressures, or metabolic trade-offs determine which defense elements are constitutively active and indispensable. Deepening the investigation into these context-dependent essentialities may uncover the evolutionary forces shaping bacterial defensome repertoires. These species-specific patterns within defense gene deployment is also consistent with experimental work performed on SRP which demonstrated that competition mechanisms are complex and mainly occur between strains, regardless of the species concern (57).

Surprisingly, phage repressors were found to be essential for most of the phage regions identified by PHASTEST, regardless of whether the region encoded a putative functional phage or a phage remnant such as the tailocin dickeyocin of the *Dickeya* genus (52). This suggests that these regions require repression in TSB 50% in the absence of a known stressor. Such tight repression is probably necessary to prevent the lytic cycle from starting. Dickeyocin production is induced by various stress such as mitomycin C, reactive oxygen species and antibiotics (52, 58). By contrast, previous research on the two complete prophages of *P. atrosepticum* SCRI1043, the Mu-like ECA41 (prophage #9) and the P2-like ECA29 (prophage #8), demonstrated their resistance to chemical or physical induction (59), suggesting that an unknown induction mechanism required to induce their lytic cycle remains to be discovered. Our work also revealed a fundamental regulatory distinction between the *Dickeya* tailocins dickeyocin and the *Pectobacterium* tailocin carotovoricin (60). Carotovoricin-producing elements lack a dedicated essential repressor gene directly associated with the carotovoricin clusters (Table 2). This absence suggests that their activation mechanism diverges from the standard transcriptional repression model controlling both complete prophages and dickeyocins. Instead, carotovoricin synthesis is known to be regulated by the bacterial SOS response via the global repressor LexA (61). The essential nature of the *lexA* gene across all *Pectobacterium* strains analyzed (OMA04984) aligns with its vital role in this distinct regulatory pathway. In addition to phage repression, the presence of a functional toxin/antitoxin pair was revealed within one of the phage regions through the essentiality of the antitoxin gene. As with many moron genes carried within phages, this toxin/antitoxin pair likely contributes to bacterial strain fitness (58). In conclusion, the TNSEEK platform, which was specifically developed for this project to compare the results of Tn-seq experiments performed on bacteria from several species and genera, has proven its proficiency here. TNSEEK can also be applied to other datasets, offering unparalleled flexibility in handling any number of strains and experiments, while ensuring the complete reproducibility of results. For example, TNSEEK will be useful for comparing the essential genes of SRP required during plant infection.

## ACKNOWLEDGMENTS

We acknowledge the ANR TNPHYTO (ANR-19-CE35-0016), the sequencing and bioinformatics expertise of the I2BC High-throughput sequencing facility (https://www.i2bc.paris-saclay.fr/sequencing/ng-sequencing), supported by France Génomique (funded by the French National Program “Investissement d’Avenir” ANR-10-INBS-09) and the Genotoul platform (https://www.genotoul.fr) for their help in genomes sequencing.

